# Novel taxonomic and functional diversity of bacteria from the upper digestive tract of chicken

**DOI:** 10.1101/2023.05.10.540237

**Authors:** Bibiana Rios-Galicia, Johan Sebastian Sáenz, Timur Yergaliyev, Christoph Roth, Amélia Camarinha-Silva, Jana Seifert

**Affiliations:** Institute of Animal Science, University of Hohenheim, Emil-Wolff-Str. 6-10, 70593 Stuttgart, Germany; HoLMiR - Hohenheim Center for Livestock Microbiome Research, University of Hohenheim, Leonore-Blosser-Reisen Weg 3, 70593 Stuttgart, Germany

**Keywords:** culture collection, chicken, crop, ileum, jejunum, *Lactobacillaceae*, microbiome, novel taxa

## Abstract

Strategies to increase the production rate of chicken for human consumption alter the natural process of microbial colonisation and the nutritional performance of the animal. The lack of sufficient reference genomes limits the interpretation of sequencing data and restrain the study of complex functions. In this study, 43 strains obtained from crop, jejunum and ileum of chicken were isolated, characterised and genome analysed to observe their metabolic profiles, adaptive strategies and to serve as future references. Eight isolates represent new species that colonise the upper gut intestinal tract and present consistent adaptations that enable us to predict their ecological role, expanding our knowledge on the adaptive functions. Molecular characterisation confirmed the classification of *Clostridium anaerovisceris* sp. nov. (DSM 113860= LMG 32675), *Clostridium butanolproducens* (DSM 115076= LMG 32878), *Ligilactobacillus hohenheimius* sp. nov. (DSM 113870= LMG 32876), *Limosilactobacillus gallus* sp. nov. (DSM 113833 =LMG 32623), *Limosilactobacillus avium* sp. nov. (DSM 113849= LMG 32671)*, Limosilactobacillus pullum* sp. nov. (DSM 115077 =LMG 32877), *Limosilactobacillus digestus* sp. nov. (DSM113835= LMG 32625) and *Limosilactobacillus difficile* sp. nov. (DSM 114195= LMG 32875). Strains of *Limosilactobacillus* were found to be more abundant in the crop, while *Ligilactobacillus* dominated the ileal digesta. Isolates from crop encode a high number of glycosidases specialized in complex polysaccharides compared to strains isolated from jejunum and ileum. These results represent the first approach for the isolation and detection of bacteria from the chicken’s upper digestive tract and describe the functional diversity of bacteria inhabiting these regions, improving the potential handling of chicken microbiota with biotechnological applications.

## Introduction

Poultry production provides one of the best sources of quality protein and plays a crucial role in sustaining livelihoods in developing countries. However, the constant pressure to supply the feeding demands has forced the use of genetic and husbandry strategies to improve the performance of animals and increase the production rate [1]. These strategies alter the microbial colonisation, physiology and nutrition of the animal, which increase the susceptibility to digestive impaired functionality [2]. Additionally, the strict regulations on the use of antibiotics in the last decade have driven research to further understand how the microbiota and immunity play a role in poultry health [3]. Such efforts have started to gain importance, from understanding the microbiome structure and dynamics to manipulating such microbes to improve chicken health or disease susceptibility and increase production efficiency [4]. Molecular-based studies have improved our understanding of the diversity, composition, and gene content of the gut microbiota in chickens. The availability of organic carbon along the intestine, and the relatively short transition time of digesta (around 8–12 h) [5, 6] favours the colonisation of heterotrophic fermentative microorganisms with a total or partial sensitivity to oxygen. In general, a healthy chicken microbiome profile will be dominated by the phyla *Firmicutes*, *Bacteroidetes*, *Actinobacteria*, and *Proteobacteria*, and its abundance will depend on the anatomical region, bird age, diet, geography and breeding conditions among others [5–11].

The recent description of Metagenome Assembled Genomes (MAGs) representing more than 1000 novel species from public chicken gut microbiome samples and caeca samples of two commercial bird genotypes, highlights how yet unexplored is the microbial diversity of chicken [8, 12]. Although the majority of these studies have a high-throughput sequencing approach and good reproducibility, the lack of sufficient reference genomes and genes limits the interpretation of sequencing data and restrains deeper analysis of detailed functions and gene catalogues construction. Studies comparing bacteria along the different anatomical sections enhance how the anatomy of the gastrointestinal tract (GIT) and its physiology influence bacterial colonisation [13–15]. The capacity of crop to store food under constant provision, allows microorganisms to initiate digestion throughout fermentation and consequently the absorption of lactic acid and butanoic acid via organic acid diffusion by the host [16].

Furthermore, crop can play a role in chicken’s health by improving nutrient digestibility and regulating the entrance of pathogenic organisms to the digestive system [14]. The importance of the small intestine lay on the high enzymatic activity enhanced by the pancreatic discharge at the anterior jejunum. In this GIT section, the density of the microbiome increases towards the ileum and enzymes of bacterial origin such as trehalase and lactase have been detected [17]. Moreover, bacterial amino acid absorption at ileum prevents the formation of consequent toxic end-products such as putrescine, cadaverine and other cytotoxic biogenic amines that could spoil meat quality [18]. Yet, the role of crop and small intestine’s microbiome within the whole digestive process have not yet been given the same relevance.

Functional analyses have described the chicken microbiota to be capable of fermenting carbohydrates and complex polysaccharides present in the host diet, producing short-chain fatty acids, organic acids, vitamins and antimicrobial compounds (e.g., bacteriocins) [19, 20]. This knowledge has improved culturomic studies, enhancing the growing conditions after observing the ecological role of groups of microorganisms within the environment. In caeca, functional studies of cultivated bacteria have described the ecology and niche occupation redundancy of strains of *Ruminococcaceae*, *Erysipelotrichaceae* or *Enterobacteriaceae*, that converge on the production of butyrate through different metabolic pathways of fermentation (acetyl-CoA, butyrate kinase or lysine fermentation) [21]. This study also reported host-guest interactive molecules, such as hyaluronidase, heparinase, chondroitinase and mucin-desulfating sulfatase detected on *Bacteroidetes* and *Firmicutes* strains isolated from caeca samples [10, 21]. The presence and expression of these molecules can indicate the level of adaptability and specialisation within the environment. Finally, the study of accessory genes such as carbohydrate transport systems, unique genes, singleton genes, or defence systems, helps to indirectly observe the organisms’ response to the environmental dynamics and helps to reconstruct the ecosystem dynamics to estimate the success of colonisation of some groups over others [22].

Despite the anatomical differences among the digestive regions, and the key role of crop and small intestine in the digestive process, most of the aforementioned studies have been done with samples from the lower digestive regions, caeca and faeces. Therefore, studying cultivable bacteria originating from the upper regions will enrich the reference genome database and provide information about the site-specific adaptations of strains colonising such GIT sections. In this study, we successfully cultivated and characterised 43 strains obtained from the upper digestive tract of chickens. The screening, annotation and taxonomic analysis of strains originating from different regions allowed us to understand the metabolic profile and adaptive strategies of the strains. Among the collection, eight isolates represented new species and expanded our knowledge of the specialised gut chicken microbiome function.

## Methods

### Animal sampling

The study was approved by the Regierungspräsidium Tübingen, Germany (HOH50/17 TE). Animals were maintained and fed *ad libitum* with a commercial corn-based diet and housed at the Lindenhof Experimental Station of the University. Ten Lohmann brown classic laying hens (LB; n = 10) of 47 to 67-week-old and six 21-day-old broilers (LB; n = 6) were euthanized by hypoxia induction with CO_2_ and immediately decapitated. The digestive tract was clamped-closed before being dissected and placed into a reductive solution of 0.5% cysteine for transportation.

### Culture media

All culture media used in this work were prepared following Tanaka, Kawasaki & Kamagata recommendations [23]. Phosphate, protein, carbohydrate and agar solutions were prepared and autoclaved separately to avoid the formation of reactive oxygen species and ketosamines during sterilisation [24]. Afterwards, solutions were equally mixed before solidification. Carbohydrate solutions with high sugar content such as MRS and PFA were sterilised at different conditions than the rest (110°C for 30 minutes) to avoid sugar oxidation. Protein and peptone solutions were prepared using peptones from soybean plants. Such measurements were taken to improve the isolation of a greater diversity when using traditional plate-dependent cultivation.

### Bacteria isolation

Samples were taken to an anaerobic station (Don Whitley Scientific, Bingley, UK) where digesta was extracted from the digestive tube and ten times serially diluted with a sterile physiological solution (0.85% NaCl). Isolates were obtained from direct cultivation of the samples and enrichment cultures by plating 0.1 mL of dilutions 10^-4^ to 10^-7^ into Tryptic Soybean Agar (TSA), Gut Microbiota Medium (GMM), Man Rogosa Sharpe Medium (fructose 1% and maltose 1%) and Poultry-Food Mixture Medium 2% (PFA).

Dilution plates were monitored every 24 h and new colonies were collected into time batches at 48, 96, 144, 216 and 360 hours. In parallel, the enrichment cultures were obtained by inoculating 1 g of the sample into each culture media and plating 0.1 mL of the diluted enrichment broth on solid agar at the same time batches and dilutions used at the direct isolation. All isolates were serially cultured into new media plates to obtain axenic cultures. Cultures were incubated at 39°C under anaerobic conditions of a gas mixture of 85% nitrogen, 10% carbon dioxide and 5% hydrogen. After the isolation procedure, all the isolates were conserved into the correspondent liquid media with 25% of glycerol solution at -80°C.

### DNA extraction

DNA extraction was performed on fresh bacterial cultures, between 48 h and 96 h of incubation, following an enzymatic lysis protocol [25]. In brief, cells were grown to a density of 1.0 (A600nm) and washed twice with phosphate buffered saline (PBS). The cell pellet was suspended on 0.2 mL of PBS buffer and incubated at 50°C with 1 U of lysozyme and 1 U of recombinant mutanolysin (A&A Biotechnology Lot.130721) for 30 min, followed by an incubation of 30 min at 56°C with 5 µL of proteinase K. One unit of RNAse A was added to each tube and incubated at room temperature for 10 minutes. Both enzymes were part of the MagAttract HMW DNA extraction Kit (Qiagen). The tubes were then centrifuged at 15,000 rpm for one minute. The supernatant was recovered and purified using magnetic beads according to the MagAttract HMW DNA Kit manufacturer’s instructions. Quantification was performed with a fluorometric method using Qubit dsDNA HS assay (Thermo Scientific, Waltham, MA, USA) in a Qubit 4 Fluorometer (Thermo Fisher Scientific, USA). DNA samples were stored at −20°C.

### Strain identification

The whole culture collection (n=689) was first screened using ARDRA digestion of the 16S rRNA gene using the restriction enzyme MspI, according to the manufacturer’s instructions (NEB Inc., Lot. R0106). Different digestion profiles (n=160) were selected and identified by analysing 16S rRNA sequences using the primers 27f and 1492r [26]. Amplicons were compared using BLAST tool [27] and aligned to the closely related species at the non-redundant GenBank 16S ribosomal RNA database from the National Center for Biotechnology Information (NCBI). Sequences were deposited at the European Nucleotide Archive database (ENA); accession numbers are listed in Supplementary Data 1.

### Genome sequencing and processing

Among the collection, 43 genomes were sequenced using a 150-bp paired-end run from Illumina NovaSeq 6000 platform. Reads were quality controlled and decontaminated from sequencing artefacts and adapters and merged using BBtools (v.37.62) [28]. Assembling was performed using SPAdes (v3.15.0) [29] under a high-coverage isolate tag for trimmed reads. Additionally, genome sequencing of some novel taxa was carried out on a PacBio Sequel II platform (Pacific Biosciences, USA) using P6 chemistry. The genomes were assembled using Trycyler (v.0.5.3) [30], utilising post-filtered reads from PacBio and short-read corrected Illumina reads. All sequencing services were carried out by Novogene Company Ltd. (Cambridge, UK). The final genome assemblies were submitted to the ENA database under the BioProject PRJEB56193. Individual genome accession numbers are given in Supplementary Data 1.

Annotation and gene prediction was done with Prokka (v1.14.5) [31]. Further metabolic pathway analyses, motif validation, and participation in individual biogeochemical transformations annotations were mapped with METABOLIC (METabolic And BiogeOchemistry anaLyses In miCrobes) (v4.0) [32]. Both annotation strategies were used to predict the main metabolic pathways in the collection.

Proteomes were used to phylogenetically place the novel genomes based on 400 marker genes with PhyloPhlAn (v3.0.2) [33] and the tree was visualised in iTOL (v6.5.8) [34] (Figure S1 and S2). Genomes were then taxonomically classified with GTDB-Tk (v2.1.0) [35]. Further, identity parameters for taxonomy delineation, such as ANI and dDHH were calculated for novel taxonomy assignments using the software EzAAI (v1.2.1) [36] and FastANI (v1.33) (https://github.com/ParBLiSS/FastANI).

Accessory features were also annotated in the genomes. Secondary metabolite biosynthetic gene clusters were detected by mining the genomes against antiSMASH 6.0. database [37]. Antiviral systems presence was identified utilising Prokaryotic Antiviral Defence LOCator (PADLOC) [38] using HMM-based homologue searches and gene presence/absence/synteny criteria. Prophage inclusions within the genomes were detected with the PHASTER search tool [39]. Only complete phage clusters, including genes related to capsid, tail proteins, proteases, and genetic material were considered complete prophage insertions.

### Large-scale genome analyses

To evaluate the relative abundance and occurrence of the isolates within the host, the assemblies were mapped using CoverM (v0.6.1) (https://github.com/wwood/CoverM), against 106 chicken DNA samples obtained from crop, ileum and caeca from the project PRJEB60928. The obtained relative abundance per region was then used to estimate the prevalence and abundance of each strain at the three regions through all the samples.

### Metabolic characterisation

Cellular fatty acids and biochemical profile of novel taxa strains were carried out by DSMZ services, Leibniz-Institut DSMZ–Deutsche Sammlung von Mikroorganismen und Zellkulturen GmbH; Braunschweig, Germany. In total, 300 mg of frozen wet-weight biomass was used for the Microbial Identification System (MIDI Inc., version 6.1). The composition of cellular fatty acids was identified by comparison with the TSBA40 naming table.

The MicroPlate phenotypic profile Biolog GEN III was used to determine 71 different carbon source utilisation assays and 23 chemical sensitivity assays. In brief, the isolate was grown on an agar medium and then suspended in a special “gelling” inoculating fluid (IF) at the recommended cell density. The cell suspension was inoculated with 100 μL per well into the GEN III MicroPlate incubated to allow the phenotypic fingerprint to form. Increased respiration reduces the tetrazolium redox dye, giving a purple colour. Negative wells and the negative control (a well with no carbon source) remain colourless. After incubation, the phenotypic fingerprint of purple wells is compared to BIOLOG’s species library to assign an identity.

## Results

In this work, a total of 689 isolates from crop, ileum and jejunum of broilers and laying hens were obtained in pure culture, 131 isolates originated from broilers and 558 isolates from laying hens. Despite cultivation media composition and different incubation times, in several cases isolates showed high isolation redundancy (Figure S3). All isolates were screened with ARDRA-fingerprinting, resulting in 160 isolates selected for identification with 16S rRNA Sanger sequencing. Once identified, the selection of strains for further analysis (genome sequencing) was based on the exclusion of risk pathogens and the inclusion of prevalent strains with the same identity but obtained from different digestive sections. Finally, the genome of 43 different strains, including eight novel species, were sequenced and functionally annotated. The collection’s diversity and source of isolation are detailed in Supplementary Data 1.

### The upper digestive tract isolates collection

The collection comprises 11 strains obtained from crop, including the families *Clostridiaceae, Lactobacillaceae, Propionibacteriaceae* and *Streptococcaceae,* with two novel species descriptions: *Clostridium butanolproducens* and *Limosilactobacillus gallus*. Fourteen strains obtained from jejunum belonging to the families *Lactobacillaceae* and *Enterococcaceae* with the novel species description *Ligilactobacillus hohenheimius* and *Limosilactobacillus pullum*, and eighteen strains from ileum, including the families *Acutalibacteraceae*, *Clostridiaceae*, *Enterococcaceae*, *Lactobacillaceae* and *Peptostreptococcaceae* from which *Clostridium anaerovisceris*, *Limosilactobacillus avium*, *Limosilactobacillus digestus* and *Limosilactobacillus difficile* are novel species.

The generated phylogenomic tree of the selected 43 strains is shown in Figure 1. More than half of the strains within the collection (n = 33) were detected at more than 50% in at least one anatomical region of the 106 metagenome samples obtained from crop, ileum and caeca of broilers, indicating their presence in dominant communities within the chicken gut microbiome. In general, all strains belonging to the genus *Limosilactobacillus* were highly prevalent across the intestine (>60%) and more abundant in crop (>0.4%) than in ileum and caeca, while for *Ligilactobacillus*, strains were highly prevalent only in ileum and caeca and more abundant in ileum (>0.9%) than in crop, indicating an anatomical section preference. However, an exception was observed for *Ligilactobacillus salivarius*, which was highly prevalent and abundant (>95% and 1.0%, respectively) in the crop.

**Figure 1.**
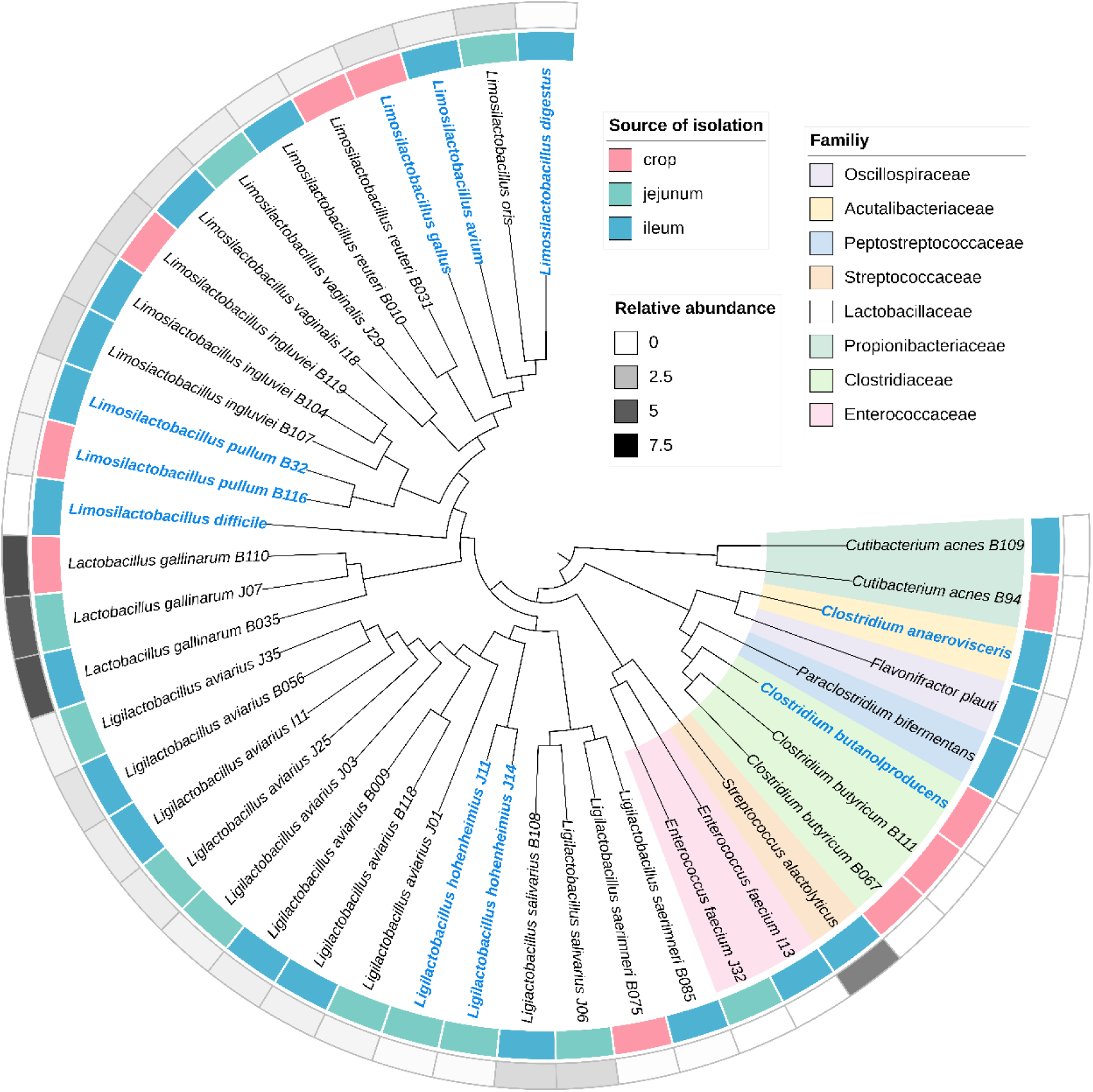
Phylogenetic tree of strains isolated from the upper digestive system of chicken based on 400 amino-acid bacterial marker sequences using PhyloPhlAn. The tree was visualized using iTol and rooted to the middle point. Novel taxa are highlighted in blue. The inner stripe depicts the source of isolation. The outside grey gradient stripe indicates the mean relative abundance of each species along gut-derived chicken DNA samples from the same chicken from where strains were obtained.

Within the collection, the species of *Lactobacillus gallinarum* and *Limosilactobacillus reuteri* were recovered from the three different sections. Both of them showed a higher abundance in the crop and ileum (>5.0 and 0.2 respectively), than in caeca (>0.3 and 0.02) and were present in all metagenomic datasets (100% prevalence) along the intestine. In addition to the abundant species, *Streptococcus alactolyticus* isolated from jejunum was highly abundant in crop (7.5%) and highly prevalent (>75%) in crop and ileum, suggesting to be a dominant member of the upper regions.

Minor abundant species *Enterococcus faecium* and *Paraclostridium bifermentans* (< 0.02%) were mapped exclusively at ileum with a prevalence of 4%. The presence of these species seems to be particularly low throughout the intestine. Contrastingly to the expected abundance of the strains at the upper regions, the species of *Flavonifractor plauti*, originally isolated from ileum was found to be abundant with 0.28% and highly prevalent at caeca (100%) compared to crop (10%) and ileum (4%). Such differences indicate a persistent species along the intestine with a higher abundance at the lower digestive parts. Finally, all *Clostridium* sp. and *Cutibacterium acnes* species were not detected in the metagenomic dataset.

### Novel taxa within the collection

Nine species represent novel taxa according to the genome-based taxonomic assignment. The taxonomic description and proposals for names are provided in the Taxonomic descriptions section. The phylogenetic tree is available at the Figures S1 and S2. Cellular fatty acids and different carbon sources utilisation were also determined. These isolates are shown in blue letters in Figure 1 and listed in Supplementary Data 1. Most of the novel taxa descriptions are based on species isolated at least twice, indicating the reproducibility of the isolation method. However, further reclassifications may occur in the future with the addition of new isolates. Among the novel taxa, six species belong to *Limosilactobacillus* genus, all isolated from crop and ileum; one to *Ligilactobacillus* genus, isolated from jejunum and two *Clostridium* species isolated from crop and ileum. (Figure 1).

The genomes of *L. gallus* and *L. hohenheimius* were already described as MAGs and given a provisional name as Candidatus *Limosilactobacillus gallistercoris* and Candidatus *Ligilactobacillus avistercoris* [40]. Since both isolates were recovered from the crop of hens and jejunum of broilers respectively, and are prevalent along the digestive tract, we propose to change the previously given names. The new assignments do not refer exclusively to faeces (suffix “stercoris”) but to the host and place of first cultivation. Hence, *L. gallus* (isolated from *Gallus gallus*) and *L. hohenheimius* (isolated at the University of Hohenheim) are proposed.

All novel species of *Lactobacillaceae* were detected on at least one of the digestive sections of crop, ileum or caeca, except for the novel species of *L. difficile* that was not detected in any sample (0% of abundance). *L. pullum* and *L. albus* isolated from crop and ileum of laying hens, had a prevalence of more than 60% and a relative abundance of more than 0.5% at crop ileum and caeca, suggesting the detection of two new key members of the chicken microbiome (Figure 1 and 2). As for *L. avium*, *L. digestus* and *L. gallus* that were prevalent with more than 50% and abundant at ∼0.05% only at crop and ileum (not at caeca). The novel species of *L. hohenheimius* was more abundant (0.1%) and prevalent (>75%) at ileum than at crop or caecum, similar to its closest related species *L. aviarius*.

**Figure 2.**
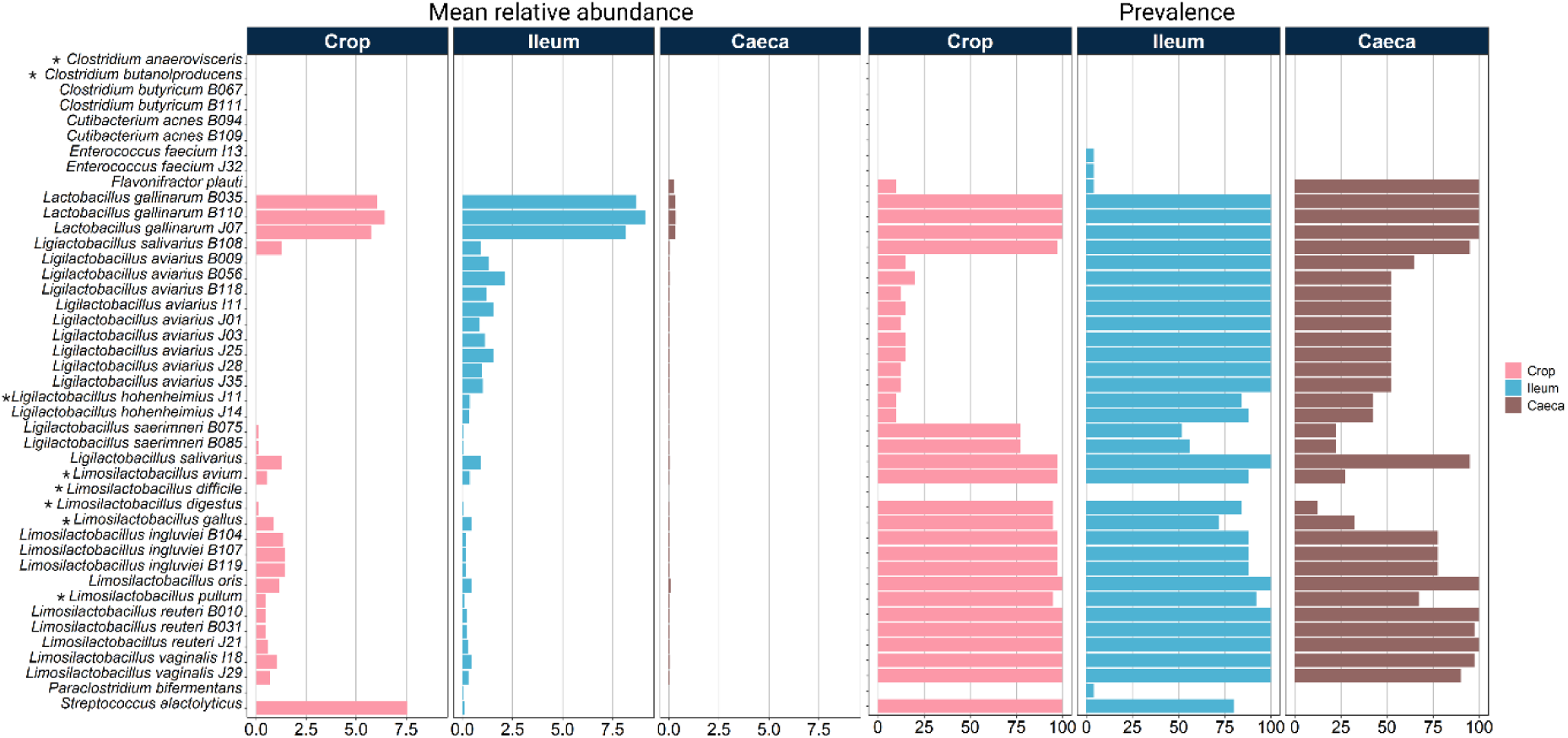
Mean relative abundance and prevalence of bacterial strains from the collection along 106 DNA digesta samples from chicken. The two panels depict the mean relative abundance and prevalence, expressed as the percentage of samples where the strain was detected at a given section. Each column represents one section: crop, ileum and caeca.

### Functional annotation

All strains within the collection, except for *Clostridiaceae* and *Oscillospiraceae* members, can produce lactic acid via lactate dehydrogenase (LDH) (Figure 3), the most common fermentative pathway, followed by the acetogenesis pathway, present in all the strains from the collection except for *Ligilactobacillus* species. Across the collection, acetogenesis was detected solely by the acetate kinase, which tells about the community’s preference for the reversible nature of the enzyme acetate kinase at the conversion and use of acetate. Additionally, this enzyme that has been reported to stimulate the chemotaxis signal system CheA-CheYBV [41], was detected in the flagellated species of our collection: *P. bifermentans, C. anaerovisceris, C. butanolproducens* and *Clostridium butyricum*.

**Figure 3.**
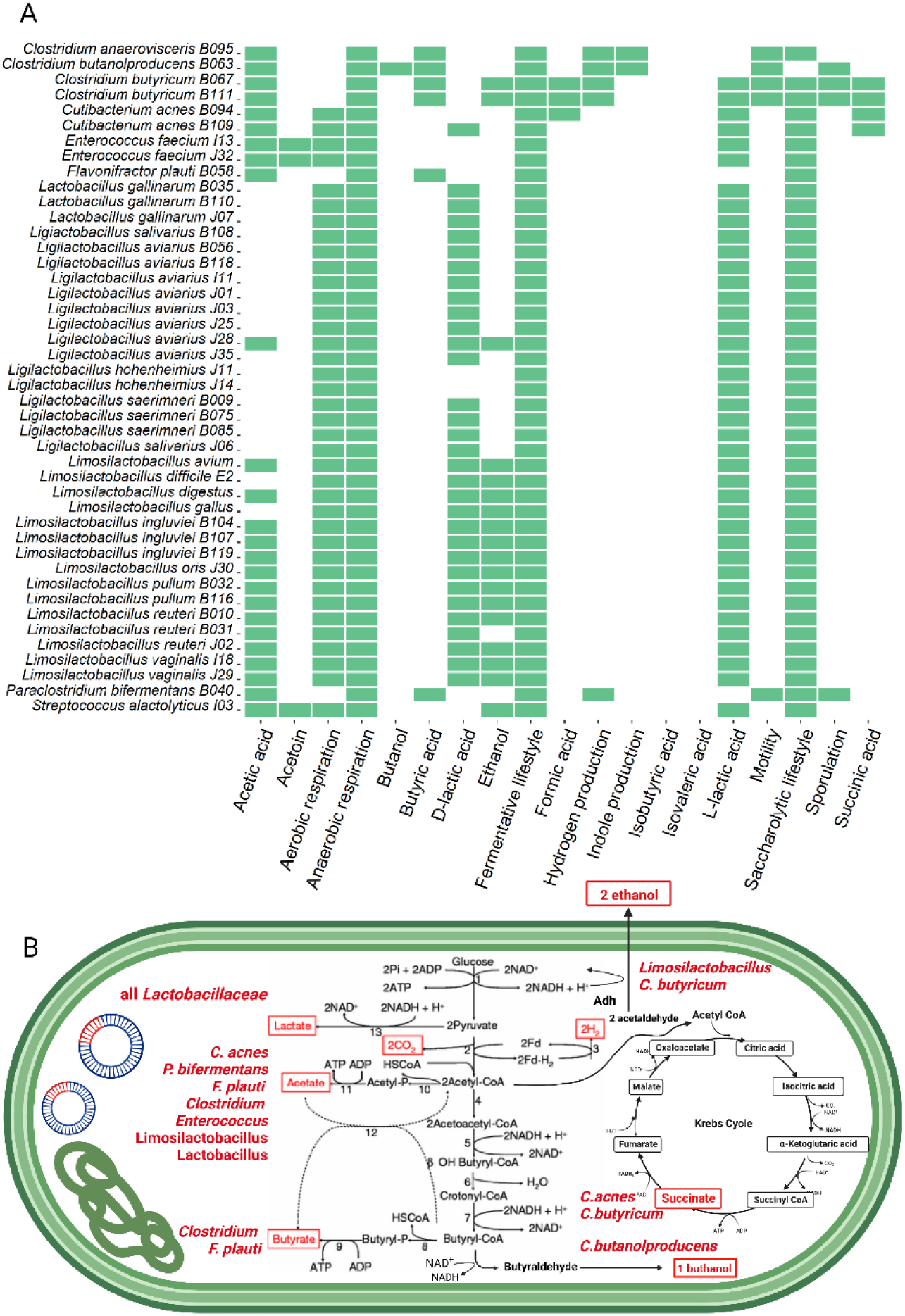
Fermentation pathways mediated by all strains from the bacterial collection of the upper digestive system of chicken. (A) Presence of fermentation pathways annotated on the genome of each strain and (B) Outline of the main pathways encoded among the bacterial collection.

Further, less common fermentative pathways mapped along the collection were butyrate fermentation, detected in *C. butyricum*, *C. anaerovisceris*, *C. acnes* and *F. plautii*, followed by succinate and formic acid fermentation, observed only in *C. butyricum* and *C. acnes* (Figure 3). All *Limosilactobacillus* and two strains of *C. butyricum* have the ability to produce ethanol from the fermentation of glucose. Only the novel taxa *C. butanolproducens* produced butanol via butanol dehydrogenase during butanoate metabolism. The scheme in Figure 4 sums up the main fermentation pathways detected on species of the collection.

**Figure 4.**
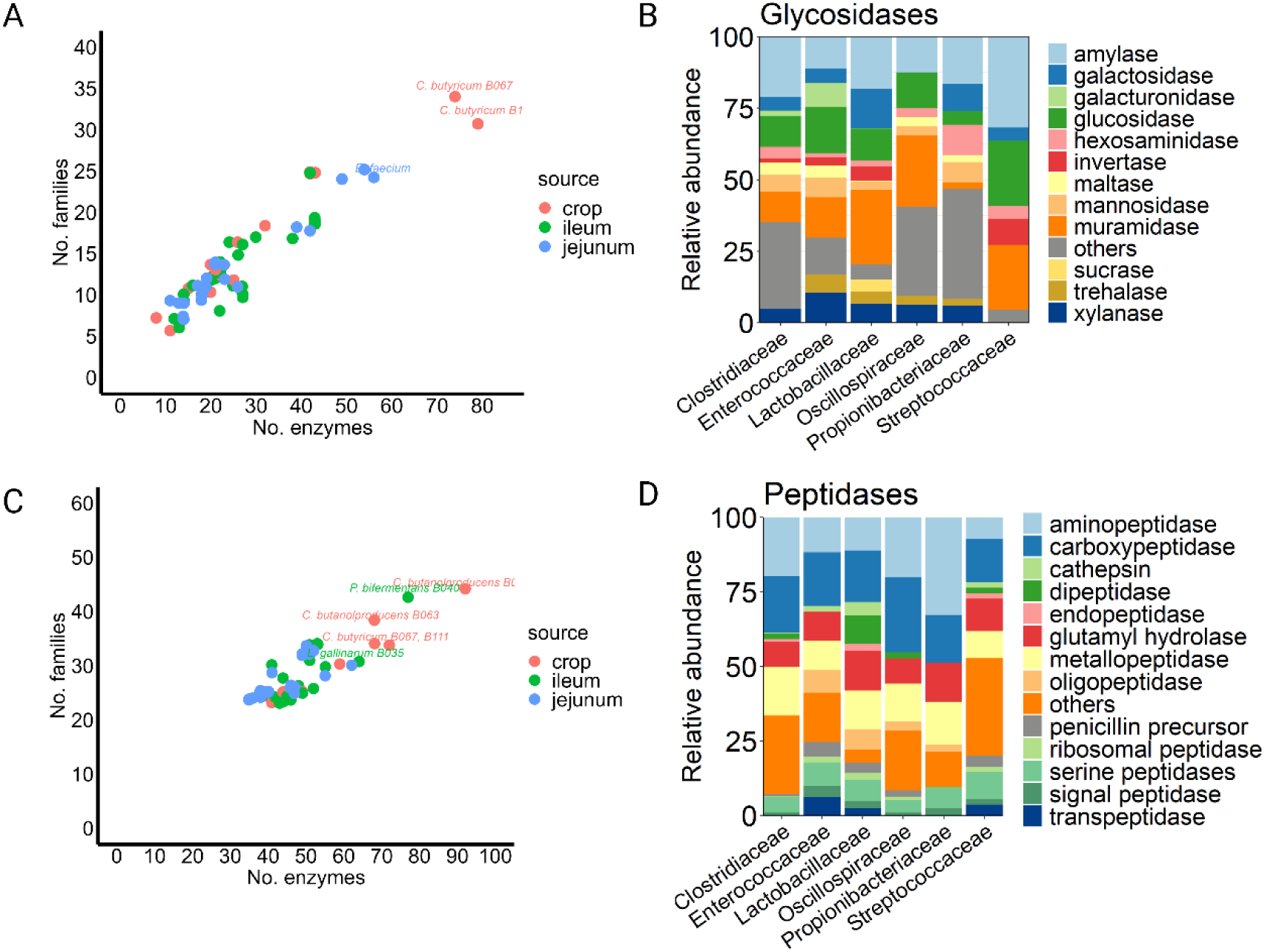
Functional enzymatic annotation across the collection of bacteria from the upper digestive system of chicken. (A and C) Total number of families of glycoside hydrolases and peptidases and (B and D) relative abundance of glycosidase and peptidases encoded by bacteria from the collection at taxonomic family level.

The following approach revealed functional differences by the number and type of genes for carbohydrate-active enzymes (CAZyme database) and peptidases (MEROPS database) throughout strains colonising different digestive sections. Despite the diverse origins of the bacterial isolates, the profile of abundant enzymes detected along the three digestive regions was conserved (Figure 4). Punctual differences were highlighted by the presence of particular taxa in a given GIT section.

Among all genes encoding glycosidic enzymes, 51% were detected in all strains and include amylases, glucosidases, muraminidases and xylanases. As for peptidase genes, 49% of signal and posttranslational peptidases and amino and carboxypeptidases were shared by all strains from the collection. Abundant genes with the respective predicted enzymatic functions are depicted in Figure 4, and their distribution among families is shown in Supplementary data 2. Commonly shared genes were higher among strains from crop and ileum (22% glycolytic and 27% peptidic) than strains from jejunum and ileum (2% glycolytic and 5% peptidic). These genes included cellobiosidases, fucosidases, galactosidases, glucosaminidases, glycogenases, hyaluronate lyases, mannosidases, peptidoglycan lyases as well as aspartyl proteases, glutamate synthases, transposases and proteasomes components found in strains from *Lactobacillaceae*, *Clostridiaceae*, *Oscillospiraceae*, *Propionibacteriaceae* and *Streptococcaceae*. Contrastingly, no shared genes for glycolytic nor peptidic enzymes were found between jejunum and crop and only one type of glucuronidase enzyme was found exclusively in jejunum strains. This means that the majority of enzymes encoded by jejunum strains might be present either in crop or ileum.

*C. butyricum* and *C. butanolproducens* isolated from crop harbour the highest counts of genes assigned to glycosidases and peptidases (Figure 3A and 3D). Strains recovered from this digestive section encode the highest proportion of section-specific glycosidases (19%) that were not detected on strains from jejunum or ileum. Such enzymes are specialised in the degradation of dextrins, furanosides, glucans, mannans and xyloglucans which underlines the potential of the local microbial community to obtain carbon from complex carbohydrate hydrolysis. Genes encoding peptidases exclusively found at crop isolated strains, were related to sporulation factors from *C. butyricum* and hydrogenase-processing endopeptidases encoded by the novel species of *C. butanolproducens* and represent 5% of the total of peptidases detected along the collection.

Strains recovered at jejunum belong either to *Enterococcaceae* or to *Lactobacillaceae* family, with *E. faecium* having the highest count and diversity of active enzymes at jejunum compared to *E. faecium* strains recovered from the ileum. Such functional diversity included chitosanase, chitinase, iduronidase, glucuronidase and hexosaminidase enzymes, all devoted to the hydrolysis of complex carbohydrates like chitin, a common polysaccharide from the exoskeleton of insects. At family level, only the group of *Enterococcaceae* presented a higher count of genes for CAZymes at jejunum compared to crop or ileum (Supplementary data 2).

As expected by the anatomy of the GIT section, ileum strains presented less genetic diversity of exclusive glycolytic enzymes than crop (5%) but higher abundance of amylases, glucosidases and muramidases, which indicates the specialisation of these bacteria to the degradation of carbohydrates previously hydrolysed at the upper parts. In this region, 14% of the exclusive peptidases found, included posttranslational peptidases, serine-protease, immune inhibitors and a protease insulin-like degrading enzyme found in the novel species *C. anaerovisceris*.

### Genes encoding host-interactive molecules

Genes for enzymatic adaptations that allow host-microbiome interactions, such as collagenases, pitrilysin and homologues of neprilysin, were detected in strains from *Clostridiaceae*, *Enterococcaceae*, *Lactobacillaceae*, *Oscillospiraceae*, *Propionibacteriaceae* and *Streptococcaceae* (Figure 5). High counts of neprilysin and pitrilysin were found in crop strains of *C. butyricum* and *C. butanolproducens*, suggesting a higher interaction of these strains to the host endothelins and glucagon/insulin hormones at regions where these strains colonise. Furthermore, collagenase producer strains were detected in higher numbers in jejunum and ileum, having *E. faecium*, *P. bifermentans* and *S. alactolyticus* the highest counts of collagenase gene clusters detected per genome.

**Figure. 5.**
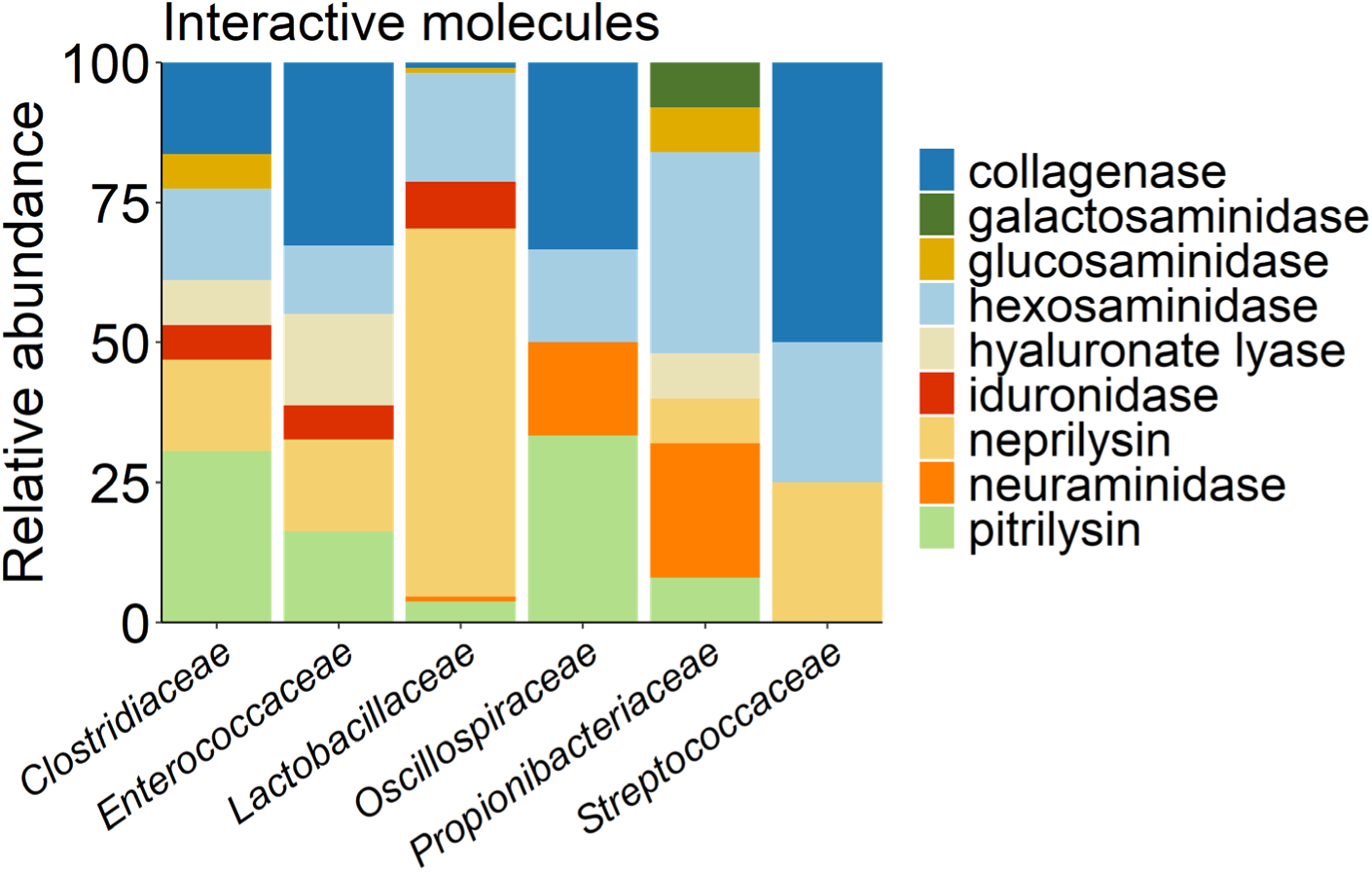
Distribution of genes encoding proteins that mediate interactions with the host across strains from the collection of bacteria from the upper digestive system of chicken grouped at family level.

Genes encoding aminidases were found mainly as hexosaminidase and sialidases at *C. acnes* and *Lactobacillaceae* strains isolated from crop, jejunum and ileum, some other less common aminidases like glucosaminidase and galactosaminidase were exclusively found at crop and ileum encoded by *F. plautii* and *C. acnes*.

The presence of hyaluronate lyase clusters along genomes of *C. butyricum*, *E. faecium* and *C. acnes* from the three sections tells about the interaction of these strains to the host digestive extracellular matrix (mucosa layer), and how they are potentially able to hydrolyse hyaluronate chains through a β–hydrolysis process. Such capability is known to be a pathogenic bacterial spreading factor either by facilitating bacterial invasion breaking the polysaccharide matrix architecture or as an antigenic disguise that prevents the recognition of bacteria by phagocytes. However, such adaptations might represent a possible specialisation to colonise mucosa.

All novel species in this work encoded gene clusters with various (de-)phosphorylation activities including protein phosphatases, inorganic phosphatases, nucleotidases, sugar phosphatases and signalling phosphatases involved in chemotaxis response. Some of which might potentially provide a source of available phosphate to the host (Figure 6).

**Figure 6.**
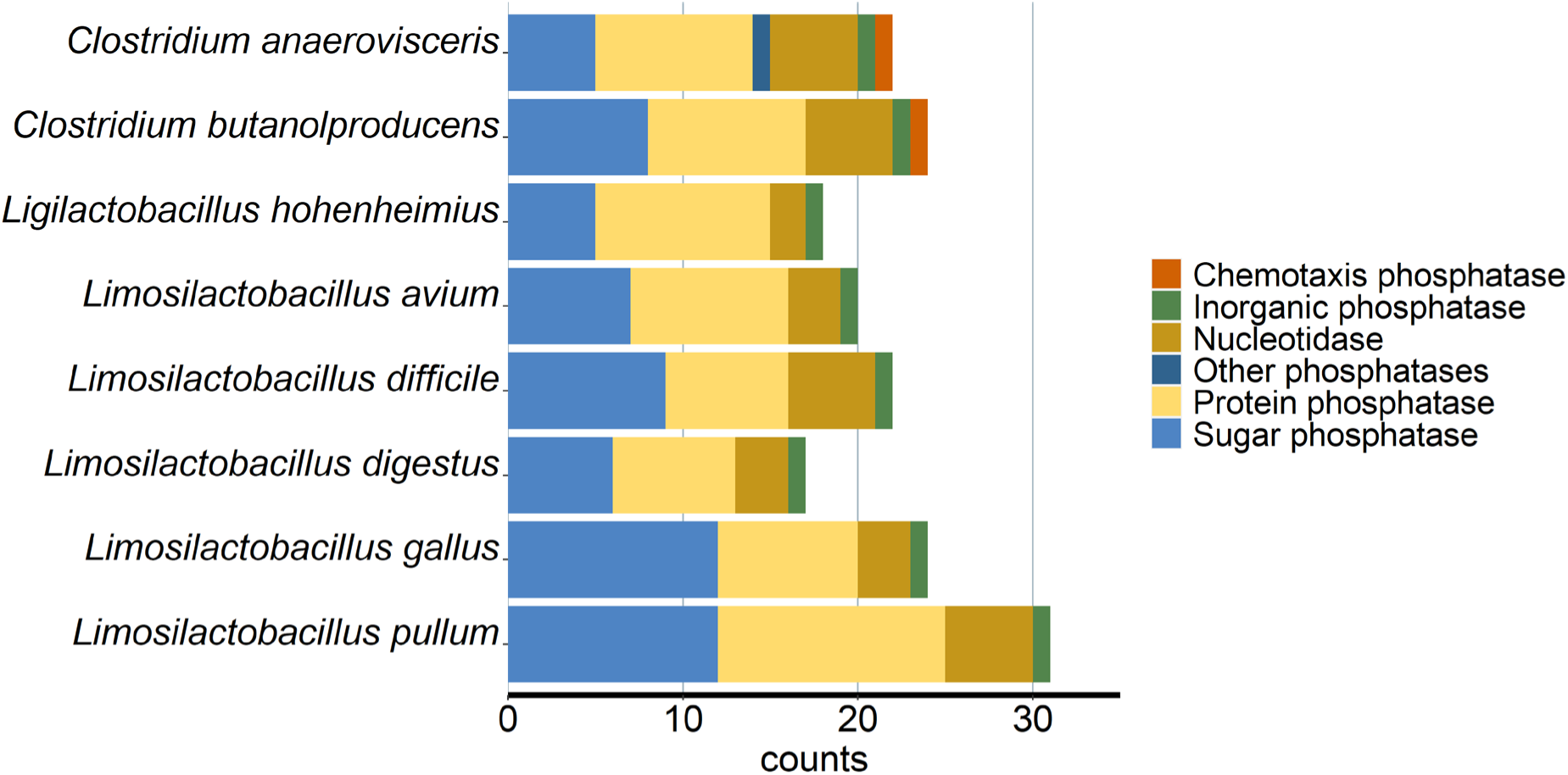
Number of phosphatases group genes present in the novel species of bacteria from the collection of bacteria from the upper digestive system of chicken.

### Accessory annotation

Beyond functional annotation, bacteria develop additional genome adaptations that reflect the complexity of interactions within members of the community and the environment (host) that assure their individual persistence in the community. The presence and transference of such features and their study, contribute to a better understanding of the bacterial community dynamics, especially in a high transitional environment such as the chicken upper digestive system. In this case, antiviral mechanisms, prophage inclusions, second metabolites detection and antimicrobial resistance were explored using accessory annotation (Figure 7). Defence systems to viral infections such as the most common restriction-modification systems were found in strains of *Limosilactobacillus* and *Ligilactobacillus*; whereas the bacterial suicide programming was encoded in all the members of the collection except for all the species of *Clostridium*, *C. acnes*, *L. hohenheimius*, *L. difficile* and *Limosilactobacillus oris*. Genes for toxin-antitoxin mechanisms (TA) were found in *Enterococcus*, *Lactobacillus* and *Ligilactobacillus*. Strains from ileum presented a higher number of antiviral defence mechanisms compared to their crop or jejunum homologues (Figure 7A). Most of these processes involve nonspecific mechanisms that target a broader range of viral invasions by blocking the infection cycle via nucleases, toxins, abortive mechanisms or the interruption of viral replication/transcription. Contrastingly, few complete prophage insertions were found in *C. butyricum*, *L. aviarius*, *Ligilactobacillus saerimneri*, *L. albus*, *L. difficile*, *L. reuteri* and *S. alactolyticus,* isolated mainly at ileum and crop (Figure 7B). Analyses of genes involved in the potential production of antimicrobials and secondary metabolites revealed one common trait found in all the strains of *Lactobacillaceae*, the type III polyketide synthase (PKSs) gene mvaS that codifies a catalytic enzyme involved in the biosynthesis of polyketides. This enzyme called hydroxymethylglutaryl-CoA synthase, is involved in the catabolism of fatty acids and enters the mevalonate pathway producing a precursor on the biosynthesis of isoprenoid compounds with the potential to inhibit pathogenic microorganisms [42]. In addition, a common antimicrobial trait detected in *Clostridium* and *Flavonifractor* strains was the cluster for ranthipeptides related to the synthesis of biologically active molecules with antimicrobial activity [43]. Very few drug resistance mechanisms were found along the collection and most of them included the genes for drug efflux transporters ABC, EmrAB, MepA, MdlAB or PatA, and the gyrase-protecting protein that confers resistance against quinolones, present on the species of *Clostridium*, *Limosilactobacillus*, *P. bifermentans* and *S. alactolyticus*. Interestingly, the ABC multidrug efflux pump EfrAB was detected in all the strains from the collection, suggesting it to be a useful trait within this environment (Supplementary data 2).

**Figure 7.**
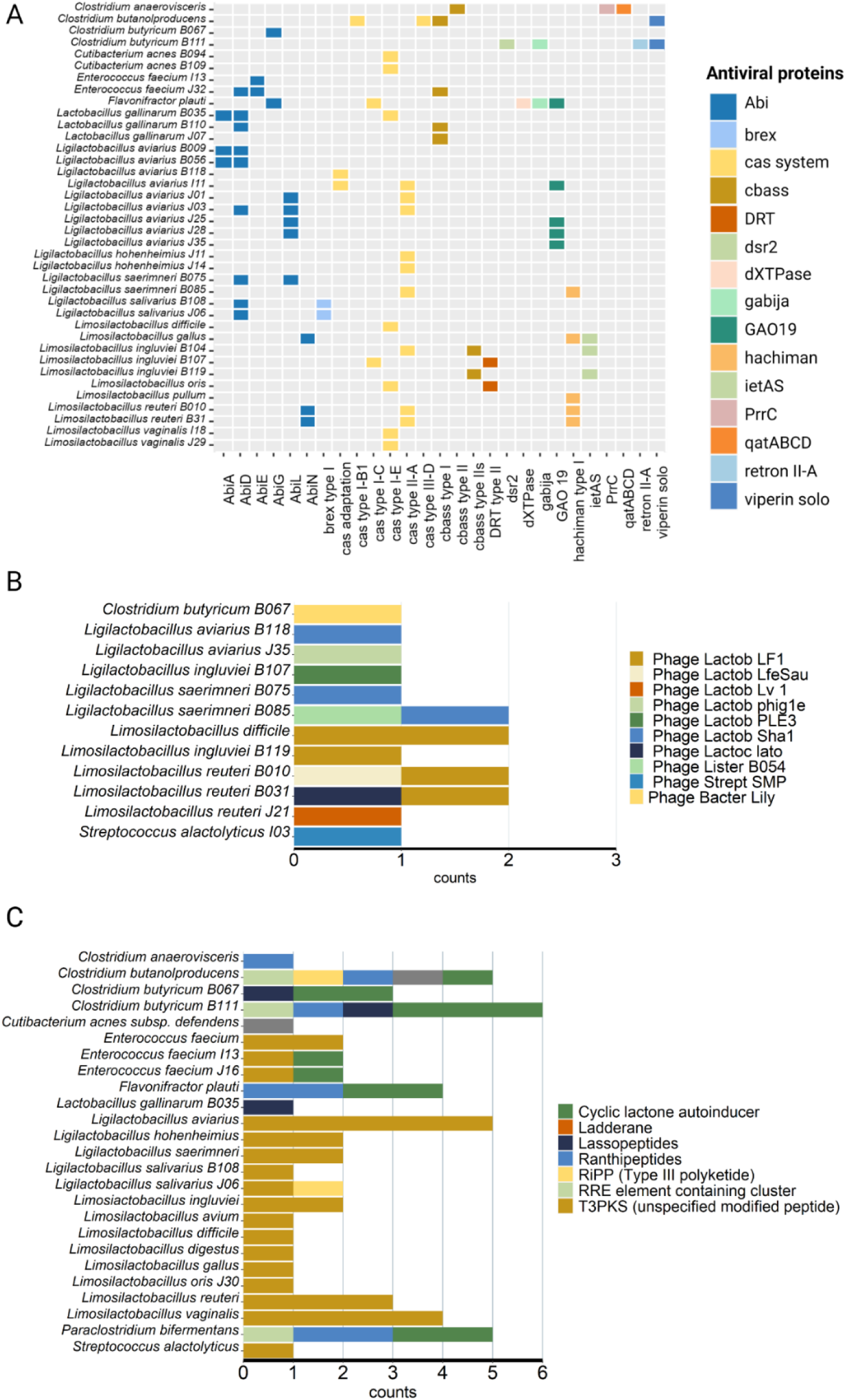
Accessory annotation for (A) antiviral defence systems, (B) prophage insertions and (C) secondary metabolite precursors detected along the upper digestive system bacterial collection of chicken.

### Taxonomic descriptions

The classification of the isolates was done based on genome sequences using the GTDB database (R207_v2) with the tool GTDB-Tk and constructing phylogenomic trees to either confirm or deny the novel delineations. Thresholds of 16S rRNA gene sequence identities were in most cases not considered as indication of a novel species. However, dDDH values <70% and ANI values <95% were considered as thresholds for separated species. Additionally, differences within-species in the G+C content of DNA (>1%) also supported the status of distinct species although not in all cases (Supplementary Data 3).

Description of *Limosilactobacillus gallus* sp. nov.: from the Latin word *gallus* (rooster, cock, male chicken) isolated in GMM from crop of brown Lohmann laying hens. The genome size is 1,861,581 bp and the mol percentage of G+C content is 50.4. The closest phylogenetic neighbour within the genus *Limosilactobacillus* (family *Lactobacillaceae*) that shares a 16S rRNA gene sequence identity of 98.92% is *Limosilactobacillus pontis*. The index dDDH and ANI of 49.2% and 83.25% respectively, and the difference in G+C content of 3.02% confirm the status of separate species. Cells are anaerobic facultative, stain positive to Gram stain and have a short rod shape. The genome encodes for aminotransferase class I and II, the Complex V (ATPsynthase), utilise lactate, cellulose and oligosaccharides as possible carbon sources and produces ethanol and lactic acid after fermentation of sugars and produces glutaredoxin. Present sugar phosphatases YbiV, YbjI and YidA, dITP/XTP pyrophosphatase, oligoribonuclease and PAP phosphatases, pyrophosphatase PpaX, phosphoserine phosphatase, tyrosine phosphatase, phosphoglycolate phosphatase, threonine phosphatase, exopolyphosphatase. The biochemical test reported that on Biolog agar media with 5% sheep blood incubated at 37°C under microaerophilic conditions, the species consumed D-maltose, sucrose, D-raffinose, fructose, lactic acid, serine, alpha keto-glutaric acid, acetic acid, butyric acid, sodium butyrate and acetoacetic acid, and tolerate minocycline, tetrazolium violet, tetrazolium blue potassium tellurite, Tween 40, 8% of NaCl and sodium bromate. The predominant cellular fatty acids are C16:0 (38.05%), C19:0 cyclo w8c (21.61%), C18:0 (17.54%), and C18:1 ω7c and C18:1 w7c (13.45%). Other fatty acids are 16:1 w6c/16:1 w7c and 16:1 w6c/16:1 w7c (4.85%). The type strain number is DSM 113833.

Description of *Limosilactobacillus avium* sp. nov.: from the Latin word *avium*, inflection form of avis (bird) isolated in MRS medium from ileum of brown Lohmann laying hens. The genome size is 2,158,112 bp and the mol percentage of G+C content is 48.5. The closest phylogenetic neighbour within the genus *Limosilactobacillus* (family *Lactobacillaceae*) that shares a 16S rRNA gene sequence identity of 99.93% is *Limosilactobacillus panis*. Although the percentage of ANI is 92.86% and the difference in G+C content is 0.41%, the index dDDH of 68.5% is under the limit of species which confirms the status of separate species. Cells are anaerobic facultative, stain positive to Gram stain and have a short rod shape. The genome encodes for flavin prenyltransferase, aminotransferase class I and II, the Complex V (ATPsynthase) and the subunit I of cytochrome ubiquinol oxidase. Utilize lactate, cellulose, rhamnosides and xylosides from hemicellulose and oligosaccharides as possible carbon substrates and produces ethanol, acetate and lactic acid after fermentation of sugars. Present phosphatases sugar phosphatases YbiV, YbjI and YidA, dITP/XTP pyrophosphatase, oligoribonuclease and PAP phosphatases, pyrophosphatase PpaX, phosphoserine phosphatase, tyrosine phosphatase, phosphoglycolate phosphatase, threonine phosphatase, exopolyphosphatase. The biochemical profile test reported that on Biolog agar media with 5% sheep blood incubated at 37°C under microaerophilic conditions, the species consumed D-trehalose, gentiobiose, glucose, D-raffinose, fructose, lactic acid, serine, sorbitol, glucoronamide, alpha keto-glutaric acid, alpha keto-butyric acid, acetic acid, butyric acid, propionic acid, formic acid, acetoacetic acid, sodium butyrate, sodium lactate and tolerate rifamycin, minocycline, potassium tellurite, guanidine HCl, vancomycin, tetrazolium, nalidixic acid, 8% of NaCl and sodium bromate. The predominant cellular fatty acids are C16:0 (52.09%), un 18.846/19:1 w6c (15.11%), 19:0 cyclo w10c/19w6 (15.11%), and C18:0 (10.97%). Other fatty acids are 18:1 w9c (5.74%), 16:1 w7c/16:1 w6c (5.27%), 16:1 w7c/16:1 w6c (5.27%) and C12:0 (4.83%). The type strain number is DSM 113849.

Description of *Limosilactobacillus pullum* sp. nov.: from the Latin word *pullum* (chicken) isolated in PFA medium from the ileum of brown Lohmann laying hens. The genome size is 1,924,125 bp and the mol percentage of G+C content is 45.9. The closest phylogenetic neighbour within the genus *Limosilactobacillus* (family *Lactobacillaceae*) that shares a 16S rRNA gene sequence identity of 99.7% is *Limosilactobacillus mucosae*. The index dDDH and ANI of 60.1% and 86.82%, respectively and the difference in G+C content is 0.53%, confirming the status of separate species. Cells are anaerobic facultative, stain positive to Gram stain and have a small rod shape. The genome encodes for a phosphoserine and the aromatic aminotransferases for histidinol, aspartate and tyrosine aromatic aminotransferase, aminotransferase class I and II, the Complex V (ATPsynthase) and the subunit I of cytochrome ubiquinol oxidase. Utilise lactate, cellulose and oligosaccharides through beta-glucosidases and beta-galactosidases as possible carbon substrates. Produces ethanol acetate and lactic acid after fermentation of sugars. Presents acylphosphatases, alkaline phosphatase, sugar phosphatases YbiV, YbjI and YidA, dITP/XTP pyrophosphatase, oligoribonucleases, pyrophosphatase PpaX, phosphoserine phosphatase, tyrosine phosphatase, phosphoglycolate phosphatase, threonine phosphatase and exopolyphosphatase. The biochemical profile test reported that on Biolog agar media with 5% sheep blood incubated at 37°C for 48h under anaerobic conditions, the species consumed dextrin, D-glucose, D-fructose, D-galactose, D-glucuronic acid, D-maltose, D-turanose, fucose, gentiobiose, glucuronamide, inosine, pectin, sucrose, stachyose, sodium lactate, sodium butyrate, D-serine, L-arginine and tolerate fusidic acid, lincomycin, minocycline, nalidixic acid, potassium tellurite, rifamycin, tetrazolium, troleandomycin, tween 40, vancomycin, 1 and 4% of NaCl. The predominant cellular fatty acids are C16:0 (41.26%), C18:1 w7c (24.53%), C18:0 (13.99%), and C19:0 cyclo w8c (10.51%). Other fatty acids are C16:1 w7c/16:1 w6c (3.34%), C14:0 (2:59%), 16:1 w7c/16:1 w6c (5.27%) and C12:0 (1.38%). The type strain number is DSM115077.

Description of *Clostridium butanolproducens* sp. nov.: from the Latin *producens* (the butanol producer) isolated in TSA medium from the crop of brown Lohmann laying hens. The genome size is 3,507,657 bp and the mol percentage of G+C content is 28.3. The closest phylogenetic neighbour within the genus *Clostridium* (family *Clostridiaceae*) that shares a 16S rRNA gene sequence identity of 98.77% is *Clostridium argentinense*. The index dDDH and ANI of 41% and 87.39% respectively confirm the status of separate species. The difference in G+C content is 0.44%. Cells are strictly anaerobic, stain positive to Gram stain and have a rod shape with endospore production. The genome encodes for branched-chain amino acid aminotransferases, aspartate and tyrosine aminotransferases, acyl-CoA dehydrogenase, formate C-acetyltransferase and the Wood Ljungdahl pathway. Present gene cluster for nitrogen fixation (nifDK), sulphur oxidation (dsrAB), sulphite reduction (asrABC), a cytoplasmic Fe-Fe hydrogenase and the ATPase proton pump type V and F (Complex V). Degrade chitin and produce ethanol butanol and acetate. Present nucleoside triphosphatases, phosphoglycolate phosphatase, dTTP/UTP pyrophosphatases, phosphosulfolactate phosphatase, arginine-phosphatases, histidinol phosphatases, CheY-P phosphatase (chemotaxis), sugar phosphatases YcdXY and YidA, dITP/XTP pyrophosphatase, and pyrophosphatases PpaX. The biochemical profile test reported that on Biolog agar media with 5% sheep blood incubated at 37°C for 47.25 h under anaerobic conditions, the species consumed dextrin, D-galacturonic acid, D-glucose, D-glucuronic acid, D-fructose, fucose, glucuronamide, L-galactonic acid lactone, L-rhamnose, pectin, acetoacetic acid, aminobutyric acid, hydroxy-butyric acid, keto-butyric acid, keto-glutaric acid, p-hydroxy-phenylacetic acid, sodium lactate, sodium butyrate, butyric acid and D-serine. It tolerates aztreonam, fusidic acid, gentiobiose, guanidine HCl, lincomycin, lithium chloride, minocycline, nalidixic acid, niaproof 4, potassium tellurite, rifamycin, tetrazolium, troleandomycin, tween 40, vancomycin and NaCl at 1, 4 and 8% concentration. The predominant cellular fatty acids are C16:0 (31.25%), C17:1 iso I/anteiso B (12.28%), C14:0 (10.77%) and C15:1 iso H/13:0 3OH (8.99%). Other fatty acids are C15:0 iso (5.26%), C18:0 (4.89 %), C 14:0 iso 3OH (3.38%), C12:0 (3.12%), C 13:0 iso (2.90%) and C 13:1 at 12-13 (2.87%). The type strain number is DSM 115076.

Description of *Clostridium anaerovisceris* sp. nov.: Latin *anaero* = no oxygen, and *visceris*= inflexion of viscus (genitive singular) that refers to any internal organ of the body (anaerobic from the internal organs), isolated in PSM medium from the ileum of brown Lohmann laying hens. The genome size is 2,921,858 bp, and the mol percentage of G+C content is 50.4. The closest phylogenetic neighbour within the family *Acutalibacteraceae* that shares a 16S rRNA gene sequence identity of 98.41% is *Clostridium* sp. Marseille-P4642. The index dDDH and ANI of 39.9% and 85.44%, respectively and the difference in G+C content of 1.51% confirm the status of separate species. Cells are strictly anaerobic, stain positive to Gram stain and have a rod shape without the production of endospores. The genome encodes for aminotransferase class I and II, phosphoserine aminotransferase, ornithine aminotransferase branched-chain amino acid aminotransferase, aspartate and tyrosine aromatic aminotransferase, histidinol aminotransferase, acyl-CoA dehydrogenase. Present gene cluster sulphur oxidation (dsrAB), sulfite reduction (asrABC), a cytoplasmic Fe-Fe hydrogenase and the ATPase proton pump type V and F (Complex V). Degrade cellulose, chitin and utilise ethanol butanol and lactate as carbon sources. Transform pyruvate to formate. Present histidinol phosphatases, dITP/XTP pyrophosphatase, CheY-P phosphatase (chemotaxis), Undecaprenyl-diphosphatase, pyrophosphatases PpaX, phosphoglycolate phosphatase, tyrosine-phosphatase, phosphoserine phosphatase, arginine phosphatases, phosphoglycolate phosphatase and nucleoside phosohatases. The biochemical profile test reported that on Biolog agar media with 5% sheep blood incubated at 37°C for 22 h under anaerobic conditions, the species consumed D-maltose, D-cellobiose, gentiobiose, D-turanose, D-raffinose, D-melibiose, D-glucose, D-mannose, D-fructose, D-galactose, 3-methyl glucose, D-fucose, L-fucose, L-rhamnose, sodium lactate, D-serine, D-glucose-6-po4, D-fructose-6-PO4, L-histidine, L-serine, pectin, D-galacturonic acid, L-galactonic acid lactone, D-glucuronic acid, glucuronamide, mucic acid, amino-butyric acid, β-hydroxy-D, L-butyric acid, alpha-keto-butyric acid, acetoacetic acid, propionic acid, acetic acid, formic acid and sodium butyrate. Tolerate troleandomycin, rifamycin, minocycline, lincomycin, guanidine HCl, Vancomycin, tetrazolium violet, tetrazolium blue, p-hydroxy-phenylacetic acid, methyl pyruvate, nalidixic acid, lithium chloride, potassium tellurite, aztreonam and sodium Bromate. The predominant cellular fatty acids are C14:0 (17.44%), 16:0 N alcohol (16.89%), C17:1 iso I/anteiso B (16.01%) and C12:0 (15.09%). Other fatty acids are C15:1 iso H/13:0 3OH (9.92%), C16:0 (8.91 %), C 16:1 w7c/16:1 w6c (5.79%) and C13:1 at 12-13 (4.26%). The type strain number is DSM 113860.

Description of *Limosilactobacillus digestus* sp. nov.: Latin *digestus* (perfect passive participle of digero =digest) isolated in MRS from the ileum of brown Lohmann laying hens. The genome size is 1,993,725 bp and the mol percentage of G+C content is 44.8. The closest phylogenetic neighbour within the genus *Limosilactobacillus* (family *Lactobacillaceae*) that shares a 16S rRNA gene sequence identity of 99.26% is *Limosilactobacillus panis*. The index dDDH and ANI of 24.2% and 77.61%, respectively, and the difference in G+C content of 3.30% confirm the status of separate species. Cells are anaerobic facultative, stain positive to Gram stain and have a short rod shape. The genome encodes for aminotransferase class I and II, the ATPase proton pump type F (Complex V). Utilise lactate, cellulose and xylose from hemicellulose as possible carbon substrates and produces formate, acetate, ethanol and lactic acid from the fermentation of sugars. Present exopolyphosphatases, nucleosidases, tyrosine phosphatases, uracil phosphatase, threonine and serine phosphatase, sugar phosphatases YbiV and YbjI, dITP/XTP pyrophosphatase and acylphosphatase. The biochemical profile test reported that on Biolog agar media with 5% sheep blood incubated at 37°C under microaerophilic conditions, the species consumed D-maltose, D-raffinose, D-lactose, D-glucose, ada-mannitol, D-serine, alpha keto-glutaric acid, acetoacetic acid and sodium butyrate. Tolerates minocycline, tetrazolium violet, tetrazolium blue, Tween 40, potassium tellurite and sodium bromate. The predominant cellular fatty acids are C16:0 (43.22%), C19:0 cyclo w8c (21.34%), C18:0 (17.60%), and C18:1 w7c (9.35%). Other fatty acids are 16:1 w7c/16:1 w6c (2.22%), C14:0 (1.79%) and 19:0 10-methyl (1.51%). The type strain number is DSM 113834.

Description of *Limosilactobacillus difficile* sp. nov.: Latin *difficile* from the Latin adjective *difficilis* (hard to isolate and grow) isolated in MRS from the ileum of brown Lohmann broilers. The genome size is 1,998,683 bp and the mol percentage of G+C content is 47.4. The closest phylogenetic neighbour within the genus *Limosilactobacillus* (family *Lactobacillaceae*) that shares a 16S rRNA gene sequence identity of 95.92% is *Limosilactobacillus pontis* LTH 2587. The index dDDH and ANI of 20.2% and 69.75% respectively, as well as the difference in G+C content of 6.06% confirm the status of separate species. Cells are anaerobic facultative, stain positive to Gram stain and have a short rod shape. The genome encodes for phosphoserine aminotransferase, aminotransferase class I and II, branched-chain amino acid aminotransferase, histidinol aminotransferase, the ATPase proton pump type F (Complex V), the subunit I of cytochrome ubiquinol oxidase and chlorite dismutase. Utilise lactate, cellulose, rhamnose and xylose from hemicellulose debranching and oligosaccharides as possible carbon substrates. Produces acetate, ethanol and lactic acid from the fermentation of sugars. Present tyrosine phosphatases, sugar phosphatases YidA, exopolyphosphatases, uracil phosphatase, tyrosine-phosphatase, dITP/XTP pyrophosphatase, histidinol phosphatase, phosphoglycolate phosphatase, acylphosphatase, threonine and serine phosphatase, Pyrophosphatase PpaX, and nucleosidase. The biochemical profile test reported that on Biolog agar media with 5% sheep blood incubated at 37°C for 18h under microaerophilic conditions this species can consume D-cellobiose, dextrin, D-fructose, D-fucose, D-galactose, D-galacturonic acid, D-glucose, D-glucuronic acid, D-lactose, D-maltose, D-mannose, D-melibiose, D-raffinose, S-salicin, D-serine, D-sorbitol, D-trehalose, D-turanose, glucuronamide, inosine, L-fucose, L-galactonic acid lactone, L-rhamnose, N-acetyl neuraminic acid, N-acetyl-D-galactosamine, N-acetyl-D-glucosamine, N-acetyl--D-mannosamine, pectin, stachyose, sucrose, acetic acid, acetoacetic acid, hydroxy-butyric acid, keto-butyric acid, keto-glutaric acid, methyl-D-glucoside, formic acid, propionic acid, sodium butyrate, sodium lactate.

Tolerates aztreonam, fusidic acid, gentiobiose, guanidine HCl, lincomycin, lithium chloride, minocycline, nalidixic acid, niaproof 4, potassium tellurite, rifamycin, tetrazolium, troleandomycin, tween 40 and vancomycin. The predominant cellular fatty acids are C16:0 (28.97%), C19:0 cyclo w8c (27.53%), C18:1 w7c (26.53%), and C18:0 (6.31%). Other fatty acids are C16:1 w7c/16:1 w6c (3.73%), C20:1 w7c (1.85) and C14:0 (1.49%). The type strain number is DSM 114195.

Description of *Ligilactobacillus hohenheimius* sp. nov.: Latin *hohenheimius* from the onomastic latinisation of Hohenheim (Hohenheim, Germany; the place of isolation). These strains were isolated in MRS from the jejunum of brown Lohmann broilers. The genome size is 1,380,885 bp and the mol percentage of G+C content is 50.5. The closest phylogenetic neighbour within the genus *Ligilactobacillus* (family *Lactobacillaceae*) that shares a 16S rRNA gene sequence identity of 93.30% is *Ligilactobacillus aviarius* NBRC 102162. The index dDDH and ANI of 13.3% and 69.45%, respectively, as well as the difference in G+C content of 10.34% confirm the status of separate species and lies in the boundary of a novel genus, however, since it does not accomplish the monophyletic ancestor requirement, it still cannot be declared a novel genus. Cells are anaerobic facultative, stain positive to Gram stain and have a short rod shape. The genome encodes for beta-glucoronidases, beta-galactosidases and hexosaminidases and the ATPase proton pump type F (Complex V). Utilise lactate and cellulose. Produces acetate, ethanol and L-lactic acid from the fermentation of sugars. Present acylphosphatase, nucleosidase, dITP/XTP pyrophosphatase, threonine and serine phosphatase, exopolyphosphatases, pyrophosphatase PpaX, tyrosine phosphatases, tyrosine-phosphatase, phosphoglycolate phosphatase. The biochemical profile test reported that on Biolog agar media with 5% sheep blood incubated at 37°C for 48 h under anaerobic conditions, the species consumed D-maltose, D-serine, D-fructose-6-PO4, L-histidine, glucuronamide, sodium lactate, sodium butyrate, quinic and mucic acid. Tolerate 8% of NaCl, minocycline, guanidine HCl, lithium chloride, potassium tellurite, Tween 40, tetrazolium violet and tetrazolium blue. The predominant cellular fatty acids are C18:0 (32.63%), C16:0 (18.81%), C19:0 cyclo w8c (11:59%), and c20:1 w7c (10.93%). Other fatty acids are C18:1 w7c (8.67%), C20:0 (6:14), C18:1 w9c (3.93%) and C14:0 (2.05%). The type strain number is DSM 113850.

## Discussion

The anatomical and physiological differences along the GIT of chickens have biased the study of gut microbiomes as independent environments with poor-traceable approaches. The upper digestive parts play a key role in the efficient utilisation of nutrients. It also represents the first defence line against pathogenic organisms that enter the digestive system with both the adaptive-innate immune functions and the presence of *Lactobacilli* capable of antagonising these organisms [44]. Yet, cultivation of bacteria from the chicken digestive system has focused on caeca and faeces samples and the search for pathogenic strains, leaving the upper regions less explored. Although cultivation methods carry a selection bias to microorganisms that adapt better to the culture conditions, and their prevalence and abundance do not always reflect the ecology and dynamics of the community, efforts in cultivating chicken bacteria help to broaden the pool of chicken gut microbiomes and manipulate strains formerly represented only by metagenomic sequencing. In this work we succeeded in isolating two *Candidate* novel species described only as metagenomics assemblies [40].

Former studies have described *Firmicutes* as a major representative of dominant groups at the crop, gizzard and small intestine [9, 45–47]. The initial screening of our isolates with ARDRA revealed that 80% belonged to Firmicutes and, among them, 20% to the *Lactobacillaceae* family. This information, together with the knowledge provided by the strain genome sequences, improves the study of community dynamics through the eyes of ecology, providing an extent of the niche role within the community even though all representatives are not yet cultured.

### The ecology of the collection along the digestive tract

The collection is composed of heterotrophic fermentative and anaerobic bacteria, that obtain energy and carbon from the oxidation of carbohydrates (mainly from glucose, pentose, maltose, cellobiose and cellulose), aliphatic amino acids (glutamine, lysine, arginine, serine) and organic acids oxidation (acetate, lactate, butyrate and long chain fatty acids). Niches might differ along the intestine depending on the availability and complexity of nutrients at the digestion stage. Still, the primary energy and carbon fluxes lead to the formation of lactate, acetate, butyrate, succinate, ethanol and butanol (Figure 3). The prevalence and relative abundance of our isolates in the hosts metagenomes, highlighted a successful isolation strategy of abundant strains within the environment. In this sense, the functional description of such genomes and their accessory genes potentially reflect their ecological role along the digestive tract.

In crop, the high substrate availability comes along with the high number of genes encoding enzymes specialised in the degradation of complex carbohydrates (dextrins, furanosids, glucans, mannans and xyloglucans) found in genomes isolated from that section. Although caecum is considered a region with complex carbohydrate fermentation [21], polysaccharide hydrolysis might be taking place in significant amounts already in the upper parts. However, due to the low absorption capacity of crop [17], substrate availability might be profited either at lower digestive regions or consumed by other commensal microorganisms. The colonisation of heterofermentative microorganisms such as *Lactobacillus* and *Limosilactobacillus* might supply the digesta content with fermentation products and simpler carbohydrates, and the detection of these microorganisms at ileum and caeca assessed by metagenome mapping, confirms the role of crop on providing metabolically active bacteria to the lower GIT sections. The dominant groups in crop were *Clostridiaceae*, *Propionibacteriaceae*, *Streptococcaceae* and *Lactobacillaceae*, specifically *Lactobacillus* and *Limosilactobacillus* genera, which dominated above *Ligilactobacilus*. Strains from these groups encoded a high number of glycolytic enzymes and antiviral mechanisms, most of which had a broad protection mechanism less specific to the target. Such metabolically active potential with less specific antiviral system profile suggests a highly transitional section, with frequent environmental changes and high niche occupation dynamics. In contrast, strains isolated from crop from the families *Lactobacillaceae*, *Clostridiaceae* and *Propionibacteriaceae*, presented the highest diversity of genes codifying antimicrobial compounds, which expression might prevent the further access of pathogenic organisms to the digestive system. Two novel taxa of *Limosilactobacillus* were cultivated for the first time. *L. gallus* isolated from the crop, and *L. pullum* obtained from crop and ileum. Both species seem to have a colonisation preference to crop but are present along the intestine in different proportions. They both presented a high count of genes encoding sugar and protein phosphatases including phosphoserine phosphatases, tyrosine-phosphatases and exopolyphosphatase, a linear polymer of residues of orthophosphate involved in energy storage, that might compete with the host for phosphate scavenging but can also be a colonisation advantage involved in bacterial motility or biofilm formation [48]. The third new species description of *C. butanolproducens* was isolated from crop at relatively high redundancy. Interestingly, this species was not found in any of the metagenomes, which can either be explained by a low abundance of the species not covered by the metagenome sequencing, a unique presence in our chickens (region-specific) or a cultivation selectivity of the isolation strategy from which these strains were obtained (See supplementary Data 1). *S. alactolyticus,* isolated initially from jejunum, depicted very high abundance at the crop, suggesting to be a dominant member of the upper regions. Although not detected in this work, *S. alactolyticus* has also been isolated from the caecum and evaluated as a potential probiotic for chicken [49].

In jejunum, the diversity of strains isolated had the least representatives regarding taxonomic diversity therefore, metabolic functionality observations were minor than at crop or ileum. Compared to the enzymatic adaptability, the amount and diversity of genes assigned to interactive molecules with the host and the accessory adaptations encoded by strains isolated in this region are the lowest of the three GIT sections. They encoded the lowest number of glycosidase families and did not possess any exclusive enzyme, except for an ɑ-glucoronidase detected in isolates of *E. faecium* from jejunum, but not present at the homologues of ileum. This enzyme would hydrolase glucuronosyl bonds in the main chain of “hardwood” xylans which might be an adaptation of *Enterococcus* strains from this section. This species was redundantly isolated in higher amounts from jejunum and ileum of broilers which suggests a high colonisation rate in this specific region. All isolates presented a cluster of bacteria involved in the secondary bile acid metabolism, an important adaptation of this region due to the high concentration of bile secretions that enter the anterior jejunum. Previous descriptions consider *E. faecium* a safe probiotic with strong intestinal adhesion and colonisation ability that can inhibit harmful microorganisms in the intestine [50]. An adhesive related protein (collagen adhesin) was found in all strains of *E. faecium*, facilitating bacterial adhesion to tissue structures with the corresponding ligand. It may also explain why *E. faecium* was not detected when mapped at other regions like crop or caeca, making it a region-specific coloniser. Studies have shown that *E. faecium* is a preventive treatment against *Salmonella* Typhimurium infection to broilers, that reduces the pathological changes in the liver and intestine and the levels of inflammatory factors such as IL-1β, TNF-α, and IL-8. Additionally, when given as an additive to chicken feed, growth performance and absorption of nutrients were significantly improved [51]. Such evidence of host and strain specificity has thriven *E. faecium,* a functional probiotic for chicken approved by the EU since 2011 [52]. An important contribution of this work is the cultivation of *L. hohenheimius*, a novel species of *Ligilactobacillus* with a very reduced genome (1.3 mbp) compared to the rest of the members of the genus that were isolated with low redundancy from the jejunum of broilers (3 isolates). The success of its isolation may be a consequence of the enrichment of the samples that preceded isolation. It had the highest abundance and prevalence at ileum, although it was also mapped at crop and caeca with lower prevalence. Functionally has a very narrow potential of partially degrading cellulose, lactose, lactate and a complete absence of amino acid utilisation genes. Such functional carbon utilisation reduction suggests an adaptation to a commensal member of the community, that depends on the nutrient supplies from the host or the environment.

The GIT section with the highest cultivation diversity and metabolic functionality in this work was ileum. Strains at this section presented the highest diversity of genes encoding peptidases and host-interactive molecules, antiviral mechanisms and prophage inclusions. The dominant group was the *Lactobacillaceae* family, specifically *Lactobacillus* and *Ligilactobacillus* genera, that were recovered in very high redundancy from both cultivation strategies (direct and enrichment), indicating anatomical dominance and colonisation preference. The highest number of novel descriptions were found in this section including *C. anaerovisceris*, not mapped at any metagenome dataset but isolated eight times under different isolation strategies; *L. avium*, *L. difficile*, *L. digestus*, and *L. pullum*, isolated with a redundancy of two to four times and detected at an abundance of ∼0.1%, a low presence compared to *Ligilactobacillus* strains. The increase of glucose absorption in the small intestine might influence substrate intake diversification to other sources of carbon such as peptides and fatty acids where these strains might find an advantage [53]. The highest number of antiviral mechanisms was found in strains of this region compared to the homologue isolates of jejunum or crop. For instance, *L. gallinarum* isolated from ileum possess the CRISPR CAS system type I-E and the abortive system AdiDL while the homologue strains of crop and jejunum presented only the cbass nuclease mechanism, indicating different adaptations to the community dynamics and higher interaction with viruses at this region. Regarding defence mechanisms against other bacteria, the type III polyketide synthase (PKSs) enzyme, detected on all the *Lactobacillaceae* family members of the collection, was related to the enzyme hydroxymethylglutaryl-CoA synthase in the synthesis of isoprenoids. However, its presence has been also associated to the lactic fermentation in fermented vegetables [54] and it is involved in the regeneration of NAD+ during lactic acid fermentation [55]. The presence of genes related to pyruvate metabolism and lactate fermentation on the collection’s genomes, suggests that hydroxymethylglutaryl-CoA synthase might interfere in both pathways, reducing NADH during lactic acid fermentation and catalysing the formation of 3-hydroxy-3-methylglutaryl-CoA (HMG-CoA) at the mevalonate pathway during the integration of secondary metabolites [42].

Genomes of all collection members included the drug resistance efflux transporter EfrAB, which provides resistance to structurally unrelated antimicrobial agents such as quinolones, tetracyclines or anthracyclines [56]. Although antibiotic resistance tests were performed only on the novel species descriptions, the presence of such a spread resistance mechanism with a broad spectrum suggests a valuable tool within the environment against different antimicrobial molecules. However, further gene expression tests must be performed to recognise this mechanism’s activity. Finally, the presence of different groups of phosphatases on all the novel species descriptions provides insights about the potential contributions of these strains to the supply of available phosphate from insoluble sources such as proteins, nucleotides, inulins, phytates and other organic sources of unavailable phosphate present in the plant-based diet given to chickens, that is not accessible to its metabolism [5, 45].

## Conclusion

This culture collection is the first bacterial repository obtained from crop and small intestine of chicken. It presents a wide enzymatic and host-interactive capacity that represents a metabolic value to the host. The redundancy of isolation from more than one source demonstrated that, although a significant portion of the genomic content is conserved, isolates hold different genomic features (e.g. carbohydrate usage, antiviral defence systems, prophage occurrence) to prosper in different environmental niches. Beyond the differences in diversity and colonisation rate through the upper digestive tract, the easiest way to observe environmental adaptations is through the comparison of similar organisms that inhabit different GIT sections such as *L. gallinarum* or *E. faecium*. This is an important contribution to better understand the adaptive response of such strains along the different regions of the digestive system. The newly obtained isolates are present along the intestine and they are consistently found within the upper GIT sections. The novel species of *Limosilactobacillus and Ligilactobacillus* represent key members of the chicken microbiome of crop or ileum and its isolation success lies in the exploration of these sections. These results represent the first deep isolation approach of bacteria from the chicken’s upper digestive system and describe the unexplored taxonomic and functional diversity of bacteria inhabiting these sections, improving the potential handling of chicken microbiota with biotechnological applications. Collectively, these findings emphasize the value of combining cultivation with metagenomics to study microbial diversity.

## Ethical approval

The use of animals in this study was reviewed and approved by the Regierungspräsidium Tübingen, Germany (approval number HOH50/17 TE).

## Acknowledgements

The authors acknowledge support by the High Performance and Cloud Computing Group at the Zentrum für Datenverarbeitung of the University of Tübingen, the state of Baden-Württemberg through bwHPC and the German Research Foundation (DFG) through grant no INST 37/935-1 FUGG. We acknowledge the kind collaboration with the culture collections at DSMZ, Braunschweig and BCCM/LMG, Gent.

